# Lesion-Level-Dependent Neuroendocrine Surge Precedes Neuroinflammation and Endoplasmic Reticulum Stress in the Hypothalamus After Spinal Cord Injury: Dual-Cohort Transcriptomic Evidence for a Temporally Ordered AVP Cascade

**DOI:** 10.64898/2026.04.19.719507

**Authors:** Lianhua Li, Hong Zeng, Ming Li, Jie Gao, Hua Chen, Bin Cai, Zhi Liu

## Abstract

**Background:** Spinal cord injury (SCI) triggers remote pathological changes in supraspinal regions, including neuroendocrine dysfunction that manifests clinically as hyponatremia and central diabetes insipidus. Clinical observations of lesion-level dependency and sequential transformation between these disorders suggest a temporally ordered hypothalamic cascade in which a compensatory arginine vasopressin (AVP)-driven neuroendocrine surge may precede a later neuroinflammation and endoplasmic reticulum (ER) stress-mediated neuronal exhaustion. Direct transcriptomic evidence for the temporal ordering of these events, however, has been lacking.

**Methods:** We performed a dual-cohort transcriptomic analysis. A discovery cohort (NCBI Sequence Read Archive PRJNA953752) comprised hypothalamic tissue from adult male Sprague-Dawley rats subjected to high-thoracic (T3) SCI, low-thoracic (T10) SCI, or sham surgery, sampled at post-injury day 7 and analyzed with edgeR/DESeq2 (|log_2_FC| > 1, P_adj_ < 0.05). An independent chronic-phase validation cohort (Gene Expression Omnibus GSE297887) of hippocampal tissue from SCI and sham mice was interrogated as a sensitive supraspinal proxy for remote neuroinflammatory and ER-stress signatures. Pre-defined gene panels covered neuroendocrine, neuroinflammation, and ER-stress/unfolded-protein-response categories.

**Results:** In the discovery cohort, high-thoracic SCI produced a lesion-level-dependent neuroendocrine surge in the hypothalamus: *Avp* (fold change 7.23; P_adj_ = 0.002), *Oxt* (fold change 14.25; P_adj_ = 2.3 × 10^−7^), and *Ucn3* (fold change 9.22; P_adj_ = 0.002) were among the most significantly upregulated genes genome-wide, whereas low-thoracic SCI failed to reach significance for any of these targets. Classical neuroinflammation markers and canonical ER-stress effectors remained transcriptionally silent (all P_adj_ > 0.69). The PERK-pathway sentinel genes *Trib3* and *Ppp1r15a/GADD34* exhibited coordinated sub-threshold trends indicative of early activation, and *Avp* expression was tightly correlated with *Mmp9* (r = 0.833; P = 0.0004). In the chronic-phase validation cohort, microglial *P2ry12* and ferroptosis signatures were significantly upregulated (*P2ry12* fold change 1.33; P = 0.008) suggesting a primed microglial state, while ER-stress effectors remained silent.

**Conclusions:** These data support a temporally ordered hypothalamic cascade after SCI in which an early compensatory neuroendocrine surge precedes — and may precipitate, through biosynthetic overload and blood–brain-barrier disruption — a subsequent neuroinflammation and ER-stress crisis. The defined molecular window between neuroendocrine activation and inflammatory/ER-stress engagement identifies a candidate therapeutic window for early neuroprotective intervention in acute SCI.

## Introduction

Spinal cord injury (SCI) is no longer regarded as a localized disruption of neural tracts but is increasingly recognized as a systemic disease that triggers widespread neuroinflammation, autonomic dysregulation, and neuroendocrine dysfunction extending far beyond the anatomical lesion.^1–4^ Transcriptomic profiling of injured central-nervous-system tissues has revealed extensive alterations in both coding and non-coding gene expression programs, underscoring the breadth and complexity of the molecular response to neurotrauma.^5^ Among the most clinically consequential yet mechanistically enigmatic remote consequences of SCI are disorders of water and sodium homeostasis — particularly hyponatremia and central diabetes insipidus (DI) — which affect up to 34.7% and 18.9% of patients with acute SCI, respectively, and are associated with hemodynamic instability, prolonged intensive care, and increased mortality.^6,7^

The clinical presentation of these neuroendocrine complications poses a diagnostic paradox. Hyponatremia and DI frequently occur sequentially within the same patient, exhibit substantial phenotypic overlap with the syndrome of inappropriate antidiuretic hormone secretion (SIADH) and cerebral salt-wasting syndrome, and display a striking lesion-level dependency — disproportionately affecting patients with cervical or high-thoracic injuries.^6–8^ Frameworks that classify these conditions as discrete, static disease entities fail to explain this dynamic clinical trajectory. A more coherent interpretation, noted in prior clinical and experimental observations, is that these syndromes represent temporally ordered stages of a single hypothalamic neuroendocrine cascade triggered by the remote effects of SCI.^6,7^

Converging lines of evidence make such a temporally ordered cascade biologically plausible. Neuroimaging meta-analyses have documented progressive gray-matter atrophy in the hypothalamus after SCI,^9,10^ and remote axonal degeneration extending into supraspinal structures has been confirmed.^11,12^ At the cellular level, microglial depletion after SCI reduces chronic brain neuroinflammation and neurodegeneration,^13^ and hypothalamic P2Y12-expressing microglia can sense hemodynamic disturbances and mount neuroinflammatory responses^14^ — a pathway that can in principle link the autonomic crisis of acute cervical SCI^15,16^ to a delayed hypothalamic inflammatory response. At the molecular level, arginine vasopressin (AVP)-producing magnocellular neurons in the paraventricular nucleus (PVN) and supraoptic nucleus (SON) are particularly vulnerable to endoplasmic reticulum (ER) stress because of their enormous protein-synthesis demands,^17,18^ and the PERK–eIF2α–ATF4–CHOP axis mediates ER-stress-induced translational arrest and apoptosis in these neurons with characteristically transient kinetics.^19–21^

Despite these converging mechanistic clues, *direct transcriptomic evidence for the temporal ordering* of these events within the SCI-affected hypothalamus is lacking. The question of temporal ordering has direct clinical implications: a defined molecular window between an initial AVP-system hyperactivation and the subsequent inflammatory/ER-stress assault would represent a candidate therapeutic opportunity for early neuroprotective intervention. Zeng et al. performed an unbiased multitissue transcriptomic analysis of the hypothalamus, spinal cord, adrenal glands, and spleen after high-thoracic (T3) and low-thoracic (T10) SCI in rats, revealing differential regulation of neuroinflammatory, Gi-GPCR signaling, circadian, and MHC-mediated immune pathways across tissues.^22^ That work established that SCI-induced immunodeficiency involves a complex neuroendocrine–immune network spanning the hypothalamic–pituitary–adrenal axis and identified tissue-specific hub genes and enriched GO pathways; however, it was designed to capture inter-tissue regulatory networks rather than intra-hypothalamic temporal dynamics, and did not specifically examine neuropeptide biosynthetic overload, ER-stress signaling cascades, or the temporal ordering of neuroendocrine versus inflammatory events within the hypothalamus — the central questions addressed here.

In the present study we conducted a dual-cohort transcriptomic re-analysis to test, at the molecular level, the prediction that an acute neuroendocrine surge in the post-SCI hypothalamus precedes — rather than accompanies — the neuroinflammatory and ER-stress cascade. Using a discovery cohort of hypothalamic RNA-sequencing data at post-injury day 7 (PRJNA953752) and an independent chronic-phase validation cohort of remote-brain RNA-sequencing data (GSE297887), we test three specific predictions of a *temporally ordered AVP cascade*: (i) an early compensatory neuroendocrine surge dominated by AVP, oxytocin (OXT), and urocortin-3 (UCN3), driven by non-osmotic stressors including neurogenic hypotension and sympathetic denervation; (ii) a transitional state characterized by sustained biosynthetic overload, blood–brain-barrier (BBB) compromise, and early PERK-pathway sentinel-gene engagement without full unfolded-protein-response (UPR) deployment; and (iii) a delayed neuroinflammatory phase, indexed by microglial activation in remote brain regions. This cascade framework serves as the scaffold for hypothesis-driven interrogation of the transcriptomic data rather than as a standalone theoretical construct.

## Materials and Methods

### Data acquisition and cohort design

A dual-cohort bioinformatic strategy was used to capture two distinct temporal windows of SCI-induced hypothalamic and remote brain pathology. The *discovery cohort* (NCBI Sequence Read Archive accession PRJNA953752) was originally generated by Zeng et al.^22^ as part of a multitissue transcriptomic study of SCI-induced immunodeficiency syndrome (SCI-IDS). The original study focused on neuroendocrine–immune regulatory networks across multiple tissues; the hypothalamic transcriptomic data have not previously been analyzed from the perspective of neuropeptide dynamics, ER-stress signaling, or the temporal cascade tested here. The cohort comprised hypothalamic tissue from adult male Sprague-Dawley rats subjected to complete spinal cord transection at the high-thoracic (T3) or low-thoracic (T10) level, or to sham surgery, with tissue harvested at post-injury day 7 (sham, n = 3; T10 SCI, n = 5; T3 SCI, n = 5). Detailed descriptions of the animal model, surgical procedures, and RNA-sequencing library preparation have been reported in the original publication.^22^ This time point falls within the predicted acute-to-subacute transition window of the cascade under test.

The *validation cohort* (Gene Expression Omnibus accession GSE297887) comprises bulk RNA-sequencing data from hippocampal tissue of adult C57BL/6J mice subjected to SCI or sham surgery (sham, n = 4; SCI, n = 3), sequenced on the Illumina NovaSeq 6000 platform (GPL24247). To our knowledge, the associated peer-reviewed publication had not appeared at the time of this submission. We acknowledge the anatomical (hippocampus vs. hypothalamus) and species (mouse vs. rat) differences between the cohorts. However, because SCI and subsequent blood-brain barrier disruption trigger a pan-encephalic (brain-wide) neuroinflammatory response driven by circulating systemic mediators, the hippocampus—a region exquisitely sensitive to such systemic insults—provides a highly validated ’surrogate window’ to index the global chronic state of supraspinal microglial activation and ER-stress signatures.

### Differential expression analysis

For the discovery cohort (PRJNA953752), raw sequence read counts were processed with a standard edgeR/DESeq2 workflow. Differential gene expression between T3 SCI versus sham and between T10 SCI versus sham was assessed using DESeq2,^23^ which models count data with the negative binomial distribution and applies Benjamini–Hochberg correction for multiple testing. Genes meeting the stringent criteria of absolute log_2_ fold change (|log_2_FC|) > 1.0 and adjusted P-value (P_adj_) < 0.05 were defined as high-confidence differentially expressed genes (DEGs). To identify potential coordinated pathway-level dynamics that might be masked by strict multiple-testing correction in a small-sample dataset, genes exhibiting an absolute log2 fold change > 1.0 but Padj ≥ 0.05 were evaluated strictly as sub-threshold exploratory trends. edgeR^24^ was used as independent verification of the primary DESeq2 analysis.

For the validation cohort (GSE297887), the pre-processed FPKM-normalized expression matrix was downloaded, containing 27,583 genes across 7 biological samples. Genes with mean FPKM < 0.1 in both groups were excluded, yielding 16,900 expressed genes. Differential expression was assessed with unpaired two-tailed Student’s *t* test with significance thresholds |fold change| > 1.5 and P < 0.05. Log_2_ fold changes were computed with a pseudocount of 0.01 to avoid division by zero.

### Targeted gene-panel analysis

To interrogate specific predictions of the cascade framework, three hypothesis-driven gene panels were pre-defined and applied across both cohorts.

#### Neuroendocrine panel

*Avp*, *Oxt*, *Ucn3*, *Asic4*, *Trh*, *Crh*, *Pdyn*, *Sst*, *Gal* — representing the core magnocellular and parvocellular secretory products of the PVN/SON.

#### Neuroinflammation panel

*Il1b*, *Tnf*, *Ccl2*, *Gfap*, *Itgam*, *Aif1*/Iba1, *P2ry12*, *Sall1*, *Hexb*, *Tmem119*, *C1qc* — covering microglial activation markers, astrocyte reactivity indicators, and classical pro-inflammatory cytokines.

#### ER-stress / UPR panel

*Hspa5*/BiP, *Atf4*, *Ddit3*/CHOP, *Xbp1*, *Eif2ak3*/PERK, *Ern1*/IRE1α, *Atf6*, *Trib3*, *Ppp1r15a*/GADD34, *Sesn2* — covering canonical UPR sensors, downstream effectors, and early sentinel genes of PERK-pathway activation.

Individual gene expression levels were compared using the appropriate test (DESeq2 P_adj_ for the discovery cohort; Student’s *t* test for the validation cohort) and visualized as bar plots with individual data points and SEM error bars.

### Cross-sample correlation and co-regulation analysis

To evaluate mechanistic coupling between the neuroendocrine surge and other pathological processes, Pearson correlation coefficients were calculated between *Avp* expression and all other genes across all 13 biological samples (sham, n = 3; T10, n = 5; T3, n = 5) from the discovery cohort. This approach exploits natural variation in injury severity across samples to identify genes whose expression scales with *Avp*, indicative of transcriptomic co-regulation. Key correlations of interest included *Mmp9* (BBB disruption), *Oxt* (magnocellular synchrony), and the broader neuropeptide family. A neuropeptide co-regulation matrix was constructed using Pearson correlation coefficients among nine selected neuropeptide genes and visualized as a lower-triangle correlogram.

### Biosynthetic-overload quantification

To estimate the ER protein-folding demand imposed by the neuroendocrine surge, cumulative neuropeptide expression (transcripts per million, TPM) was calculated for five major secretory products (*Avp*, *Oxt*, *Ucn3*, *Trh*, *Crh*) in sham versus T3 SCI groups. These neuropeptides are all synthesized as larger precursor polypeptides requiring ER-dependent folding, processing, and packaging into dense-core vesicles; their absolute expression levels therefore directly reflect the biosynthetic burden on the ER machinery of magnocellular neurons.

### Statistical analysis and visualization

Differential expression for the discovery cohort used DESeq2 (R 4.2.1) with Benjamini–Hochberg correction. Validation-cohort analyses used unpaired two-tailed Student’s *t* tests. Cross-sample correlations used Pearson’s r. Significance thresholds, pseudocounts, and filter criteria are detailed above. Significance levels are reported as ***P < 0.001; **P < 0.01; *P < 0.05; ns, not significant. All figures were exported at 300–600 dpi resolution. No adjustment was applied for the *a priori* selection of the three hypothesis-driven gene panels; this is addressed in *Limitations*.

### Data and code availability

The RNA-sequencing data analyzed here are publicly available through the NCBI Sequence Read Archive (PRJNA953752; original publication: Zeng et al.^22^) and the Gene Expression Omnibus (GSE297887; BioProject PRJNA1266676). All analysis scripts and processed data files used to generate the figures and results are available from the corresponding author on reasonable request. No new animal or human experiments were conducted for this study.

### Ethical approval

This study is a secondary analysis of publicly available transcriptomic datasets and does not involve new animal or human experiments. The discovery cohort (PRJNA953752) was generated under approval from the Animal Welfare Ethics Branch of the Shanghai Jiao Tong University School of Medicine Bioethics Committee and the Institutional Animal Care and Use Committee of Shanghai Ninth People’s Hospital (protocol SH9H-2020-T286-1), as reported in the original publication.^22^ The validation cohort (GSE297887) was deposited as a public dataset; associated ethical approval was obtained by the original depositors.

## Results

### High-thoracic SCI triggers a lesion-level-dependent neuroendocrine surge in the hypothalamus

To determine whether the transcriptomic response of the distal neuroendocrine center depends on the neurological level of injury, we analyzed the discovery cohort (PRJNA953752). Differential expression analysis with DESeq2 identified 333 genes passing the |log_2_FC| > 1 threshold in the T3 SCI versus sham comparison (266 upregulated, 67 downregulated). In the T10 SCI versus sham comparison, 124 genes passed the same fold-change criterion (66 up, 58 down), with 36 genes overlapping between the two comparisons.

Application of the stringent statistical threshold (P_adj_ < 0.05) revealed a striking pattern: only 8 genes in the entire T3 transcriptome achieved genome-wide significance after multiple-testing correction, and the four most significant were all core neuroendocrine genes. *Oxt* was the most significantly upregulated gene genome-wide (fold change 14.25; P_adj_ = 2.30 × 10^−7^), followed by *Asic4* (fold change 2.11; P_adj_ = 1.96 × 10^−5^), *Avp* (fold change 7.23; P_adj_ = 2.04 × 10^−3^), and *Ucn3* (fold change 9.22; P_adj_ = 2.04 × 10^−3^) (**Figure 1A**, **Figure 2A**). The single most robust and statistically unequivocal transcriptomic event in the hypothalamus at post-injury day 7 was therefore the upregulation of the neuroendocrine secretory apparatus.

**Figure 1.**
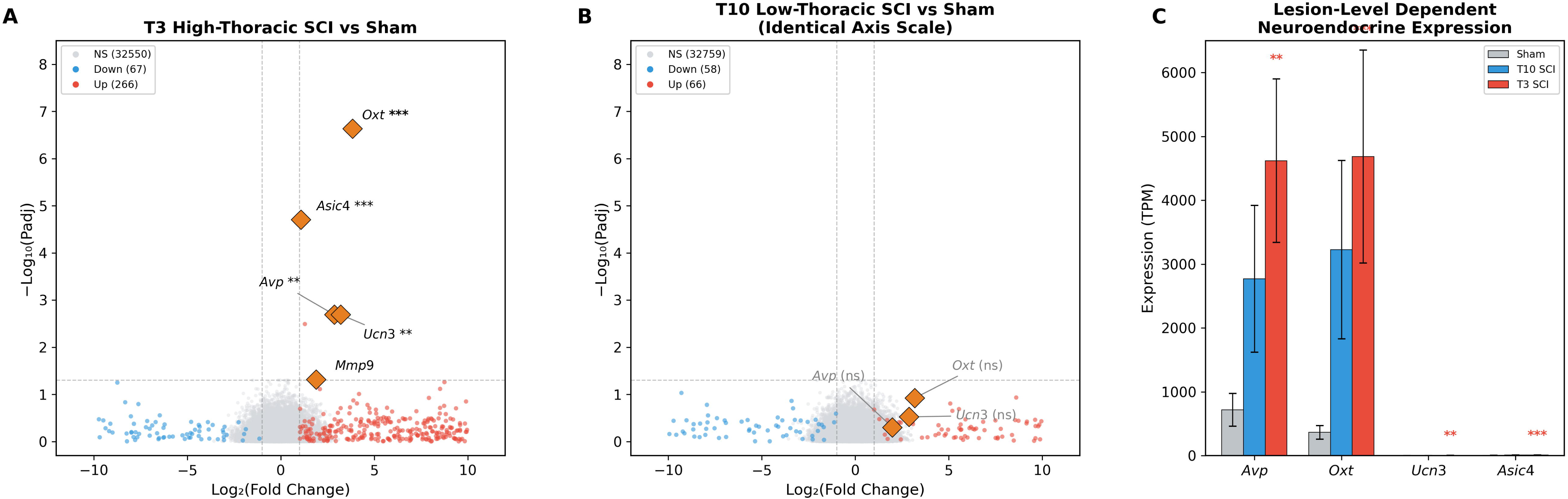
Lesion-level dependent transcriptomic reprogramming and neuroendocrine disruption in the hypothalamus following spinal cord injury (Discovery Cohort: PRJNA953752). (A) Volcano plot displaying the transcriptomic landscape of the hypothalamus following high-thoracic (T3) spinal cord injury compared to sham controls. Red and blue dots indicate significantly upregulated and downregulated genes, respectively (|Log2FC| > 1). Key neuroendocrine targets (Avp, Oxt, Ucn3, Asic4) achieving genome-wide significance (Padj < 0.05) are highlighted. (B) Volcano plot comparing low-thoracic (T10) SCI to sham controls, plotted on the identical axis scale as (A) for direct comparison. Note the striking absence of statistically significant neuroendocrine transcriptional alterations after correction for multiple testing, despite directional fold-change trends in Avp and Oxt. (C) Bar plot illustrating the absolute expression levels (Transcripts Per Million, TPM) of core neuroendocrine hub genes across the three experimental groups. Only the T3 SCI group (red) exhibits robust hyperactivation, whereas the T10 SCI group (blue) remains comparable to baseline sham levels (gray). Data are presented as mean ± SD. **Padj < 0.01; ***Padj < 0.001. Differential expression analysis was performed using edgeR/DESeq2 with Benjamini–Hochberg correction.

**Figure 2.**
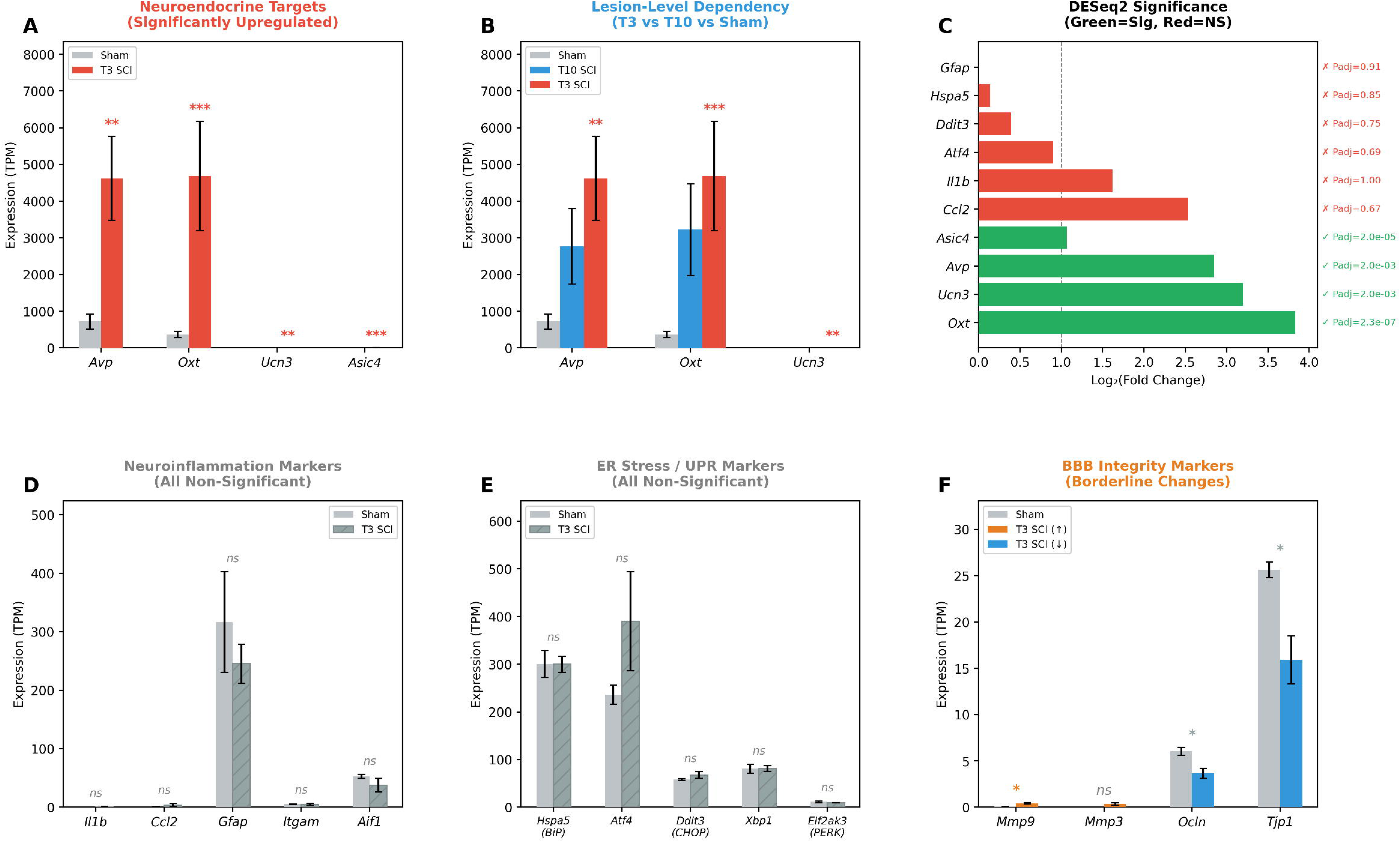
Targeted gene panel analysis reveals a neuroendocrine surge without concurrent neuroinflammation or ER stress in the post-SCI hypothalamus (Discovery Cohort: PRJNA953752, Day 7). (A) Bar plot comparing expression levels (TPM) of core neuroendocrine targets (Avp, Oxt, Ucn3, Asic4) between Sham and T3 SCI groups. All four genes achieved genome-wide significance after multiple testing correction (**Padj < 0.01; ***Padj < 0.001). (B) Lesion-level dependency comparison showing expression of Avp, Oxt, and Ucn3 across Sham (gray), T10 SCI (blue), and T3 SCI (red) groups. Statistical significance was observed exclusively in the T3 group. (C) Horizontal bar chart summarizing Log2(Fold Change) and DESeq2-corrected Padj values for neuroendocrine targets (green, significant) versus neuroinflammation and ER stress markers (red, non-significant), demonstrating the striking dissociation between robust neuroendocrine activation and transcriptional silence of inflammatory/stress pathways. (D) Expression levels of classical neuroinflammation markers (Il1b, Ccl2, Gfap, Itgam, Aif1) in Sham versus T3 SCI groups. All markers remained non-significant (ns) after correction, indicating that the hypothalamic inflammatory microenvironment has not yet developed at post-injury day 7. (E) Expression levels of canonical ER stress/unfolded protein response (UPR) effectors (Hspa5/BiP, Atf4, Ddit3/CHOP, Xbp1, Eif2ak3/PERK) in Sham versus T3 SCI groups. No significant changes were detected in any component of the UPR cascade (all ns), demonstrating that ER stress has not materialized at this time point despite maximal neuroendocrine output. (F) Expression levels of blood–brain barrier (BBB) integrity markers. Mmp9 and Mmp3 (orange, upregulated) and Ocln and Tjp1 (blue, downregulated) show borderline changes consistent with early BBB compromise. *P < 0.05 by unpaired Student’s t-test; ns, not significant. Bars represent mean ± SEM; individual data points are shown where applicable.

The lesion-level dependency of this response was confirmed by direct comparison with the T10 group. Although *Avp* (fold change 3.95), *Oxt* (fold change 9.01), and *Ucn3* (fold change 7.33) showed similar directional trends after T10 SCI, none achieved significance after correction (all P_adj_ > 0.12) (**Figure 1B**, **Figure 2B**). Absolute expression levels (TPM) further illustrated this dichotomy: AVP expression in the T3 group reached 4621 ± 2560 TPM compared with 720 ± 363 TPM in sham controls, whereas the T10 group (2771 ± 2300 TPM) showed intermediate values with high inter-animal variability (**Figure 1C**). These data establish a statistically validated lesion-level dependency that mirrors the clinical observation that dysnatremia after SCI disproportionately affects patients with high-cervical or upper-thoracic injuries.^6,7^

### Classical neuroinflammation and ER-stress pathways remain transcriptionally silent at post-injury day 7

A central prediction of the temporally ordered cascade is that the neuroendocrine compensatory surge *precedes* the neuroinflammatory/ER-stress assault. To test this, we performed targeted interrogation of the neuroinflammation and ER-stress panels in the T3 SCI hypothalamic transcriptome (**Figure 2D, E**).

#### Neuroinflammation markers

Despite the massive neuroendocrine response, classical markers of microglial activation and neuroinflammation remained transcriptionally inert. *Il1b* showed a fold-change trend (fold change 3.08) with P_adj_ = 1.00, indicating complete statistical non-significance after correction. *Gfap* (fold change 0.86; P_adj_ = 0.905) showed no change, and *Itgam*/CD11b, *Aif1*/Iba1, and *Tnf* were not among the differentially expressed genes. *P2ry12* showed a non-significant downward trend (fold change 0.52; P_adj_ = 0.665) in the hypothalamus at this time point — a finding contrasted below with its significant upregulation in the chronic-phase validation cohort.

#### ER-stress/UPR effectors

Comprehensive interrogation of the canonical ER-stress pathway revealed no significant activation at any level of the UPR cascade. The master chaperone *Hspa5*/BiP (fold change 1.11; P_adj_ = 0.855), the transcription factor *Atf4* (fold change 1.86; P_adj_ = 0.692), the pro-apoptotic effector *Ddit3*/CHOP (fold change 1.31; P_adj_ = 0.750), the splicing factor *Xbp1* (fold change 1.00), and the upstream sensor *Eif2ak3*/PERK (fold change 0.82) all remained at baseline levels (**Figure 2C, E**). This absence of UPR transcriptional activation, occurring simultaneously with extreme neuroendocrine hyperactivation, indicates that at post-injury day 7 the hypothalamus is in a state of maximal biosynthetic output without yet showing the downstream stress response predicted to follow.

### Sub-threshold trends suggest coordinated early PERK-pathway sentinel engagement

Although the canonical UPR effectors remained silent, an exploratory analysis of the sub-threshold transcriptional shifts revealed a coordinated directional trend among several ER-stress-adjacent sentinel genes consistent with the earliest stages of PERK-pathway activation (**Figure 3E**). *Trib3* (fold change 5.9; log_2_FC = 2.78; P_adj_ = 0.271) is a well-established transcriptional target of the ATF4–CHOP axis and functions as a pseudokinase that amplifies ER-stress signaling by inhibiting AKT-mediated survival pathways.^25^ While failing to reach genome-wide significance, its pronounced directional shift (log2FC = 2.78) suggests that ATF4 transcriptional activity may be functionally engaged, preceding the transcriptional surge of CHOP itself. *Ppp1r15a*/GADD34 (fold change 2.5; log_2_FC = 1.54) encodes the regulatory subunit of the phosphatase that dephosphorylates eIF2α, providing negative feedback to the PERK–eIF2α axis; its upregulation is a hallmark of PERK-pathway activation specifically documented in hypothalamic neurons under stress.^20^ This pathway-level pattern was further supported by sub-threshold trends in other stress-responsive genes, including Sesn2 (fold change 2.4), *Gadd45g* (fold change 1.8), and the metallothioneins *Mt1* (fold change 14.5) and *Mt2a* (fold change 9.05), classical indicators of oxidative stress and metal-ion dyshomeostasis. Together, these sentinel genes are consistent with a hypothalamic microenvironment transitioning from pure neuroendocrine hyperactivation toward cellular stress. Together with the inflammatory and microglial trajectories defined by the validation cohort (see below), these observations support a temporally ordered cascade model in which the neuroendocrine surge at day 7 precedes and primes both the predicted subacute ER-stress peak and the subsequent chronic microglial activation (**Figure 3F**; see **Supplementary Figure S1** for detailed mapping of phase-specific therapeutic windows).

**Figure 3.**
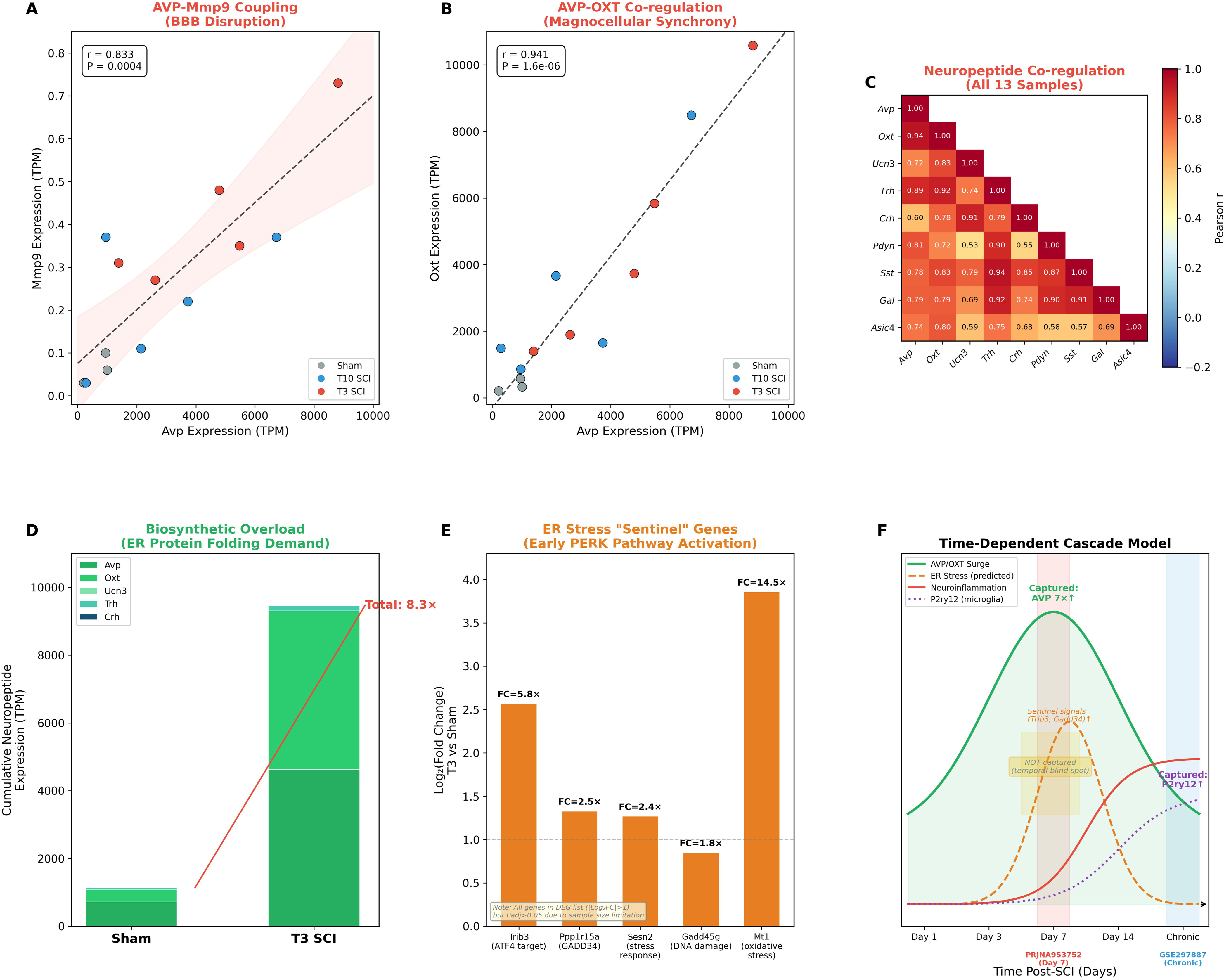
Multi-dimensional evidence linking the neuroendocrine surge to the AVP Depletion Framework (Discovery Cohort: PRJNA953752, Day 7). (A) Scatter plot demonstrating the strong positive correlation between Avp and Mmp9 expression across all 13 biological samples (Sham, n = 3; T10, n = 5; T3, n = 5). Pearson r = 0.833, P = 0.0004. The dashed line represents the linear regression fit with 95% confidence interval (shaded area). This tight coupling indicates that samples with the most intense AVP transcription also exhibit the greatest BBB enzymatic breakdown, linking the early neuroendocrine compensation with the structural prerequisite for the subsequent delayed inflammatory infiltration. (B) Scatter plot of Avp versus Oxt expression (Pearson r = 0.941, P = 1.6 × 10⁻⁶), confirming near-perfect synchronization of the magnocellular neurosecretory system and supporting the rationale for dual-receptor (AVP + OXT) replacement therapy. (C) Lower-triangle Pearson correlation matrix of nine neuropeptide genes across all 13 samples. The predominantly red heatmap (r > 0.60 for most pairs) reveals a pan-neuropeptide co-regulation network, demonstrating that the post-SCI hypothalamic response represents a global neuroendocrine mobilization rather than an AVP-specific event. (D) Stacked bar plot quantifying the cumulative neuropeptide expression burden (TPM) of five major secretory products (Avp, Oxt, Ucn3, Trh, Crh) in Sham versus T3 SCI groups. The 8.3-fold increase in aggregate neuropeptide mRNA reflects the extraordinary biosynthetic demand placed on the endoplasmic reticulum protein folding machinery of magnocellular neurons—the molecular precondition for the subsequent ER-stress cascade. (E) Bar plot of Log2(Fold Change) for ER stress ‘sentinel’ genes exhibiting sub-threshold exploratory trends: Trib3 (FC = 5.9×, a direct ATF4–CHOP transcriptional target), Ppp1r15a/GADD34 (FC = 2.5×, the PERK pathway negative feedback regulator), Sesn2 (FC = 2.4×, a stress-responsive gene), Gadd45g (FC = 1.8×, DNA damage-inducible), and Mt1 (FC = 14.5×, a metallothionein indicative of oxidative stress). Note: all depicted sentinel genes passed the |Log2FC| > 1 threshold but did not reach Padj < 0.05. They are presented here to illustrate a coordinated pathway-level trend rather than strict differential expression. Their biological identities as established PERK pathway components provide exploratory clues for early UPR engagement preceding full CHOP activation. Dashed line indicates |Log2FC| = 1. (F) Schematic of the time-dependent cascade model integrating findings from both cohorts. The x-axis represents post-injury time; colored curves depict the predicted temporal trajectories of AVP/OXT surge (green), ER stress (orange dashed, predicted transient peak), neuroinflammation (red), and P2ry12 microglial activation (purple dotted). Red and blue shaded boxes indicate the sampling windows of the discovery cohort (PRJNA953752, Day 7) and validation cohort (GSE297887, chronic phase), respectively. The yellow box highlights the ‘temporal blind spot’—the subacute window during which ER stress is predicted to peak but was not captured by either dataset. This model demonstrates that the neuroendocrine surge is already fully established at Day 7, while the inflammatory-ER stress assault materializes at later time points.

### Neuroendocrine surge is tightly coupled with BBB disruption and a pan-neuropeptide co-regulation network

To explore mechanistic relationships linking the neuroendocrine surge to other pathological processes, we performed cross-sample Pearson correlation analysis across all 13 biological samples (sham, T10, T3).

#### AVP–Mmp9 coupling

A highly significant positive correlation was identified between *Avp* and *Mmp9* expression (r = 0.833; P = 0.0004) (**Figure 3A**). MMP-9 is a matrix metalloproteinase directly responsible for the degradation of BBB tight-junction proteins and basement-membrane components.^26,27^ This tight coupling indicates that samples with the most intense AVP transcription also exhibit the greatest degree of BBB enzymatic breakdown, linking the hemodynamic crisis with the structural compromise that ultimately permits peripheral inflammatory mediators to access the hypothalamic parenchyma. This BBB compromise is further supported by the complementary upregulation of *Mmp3* and the downregulation of tight junction proteins *Ocln* and *Tjp1* **(Figure 2F).** Within the T3 group alone this correlation remained strong (r = 0.893; P = 0.041), confirming that the association is not driven solely by inter-group differences.

#### AVP–OXT magnocellular synchrony

*Avp* and *Oxt* expression were near-perfectly correlated across all samples (r = 0.941; P = 1.6 × 10^−6^) (**Figure 3B**), confirming synchronized activation of the entire magnocellular neurosecretory system, consistent with the shared SON/PVN anatomical localization and overlapping stimulus-secretion coupling of these two neurohypophyseal systems.^28,29^ Extending this pairwise synchrony to the broader neuropeptide repertoire, a lower-triangle correlation matrix of nine hypothalamic neuropeptide genes revealed a densely positive co-regulation network spanning both magnocellular and parvocellular secretory lineages (**Figure 3C**). This pattern indicates that the post-SCI hypothalamic response is not an AVP-specific event but a global neuroendocrine mobilization, consistent with a shared upstream drive acting on multiple neurosecretory cell types.

### Quantitative modeling of hypothalamic biosynthetic overload

To estimate the magnitude of the ER protein-folding demand imposed by the neuroendocrine surge, cumulative neuropeptide expression (TPM) was calculated for five major secretory products (*Avp*, *Oxt*, *Ucn3*, *Trh*, *Crh*) in sham versus T3 SCI groups (**Figure 3D**). Cumulative neuropeptide transcript abundance was substantially higher in the T3 group, indicating that the ER machinery of magnocellular neurons is being asked to process a markedly greater load of secretory-pathway cargo at day 7. This biosynthetic-overload observation is concordant with the concurrent induction of PERK-pathway sentinel genes (*Trib3*, *Ppp1r15a*/GADD34) in the same tissue.

### Chronic-phase validation cohort: microglial activation emerges while ER stress remains silent

To obtain an independent snapshot of the trajectory of neuroinflammatory and ER-stress signatures at a later time point, we interrogated the chronic-phase validation cohort (GSE297887). Given that SCI induces a pan-encephalic inflammatory response, we utilized hippocampal tissue as a sensitive supraspinal proxy to gauge the delayed neuroinflammatory state. In hippocampal tissue from SCI mice, the microglial sentinel *P2ry12* was significantly upregulated (fold change 1.33; P = 0.008), accompanied by elevated expression of homeostatic microglial markers *Sall1* (fold change 1.24; P = 0.003), *Aif1/Iba1* (fold change 1.18; P = 0.034), and *Hexb* (fold change 1.19; P = 0.044) **(Figure 4A, B).** Furthermore, pathway enrichment analysis of the upregulated genes highlighted significant alterations in myelination and, notably, the ferroptosis signaling pathway (*Alox12, Slc40a1*) **(Figure 4C).** This molecular signature suggests a non-classical, primed microglial state. Principal component analysis confirmed robust genome-wide separation between the injured and sham groups **(Figure 4D)**. *In contrast, canonical ER-stress effectors* remained transcriptionally silent (*Ddit3*/CHOP fold change 0.99, P = 0.903; *Atf4* fold change 1.05, P = 0.499; *Eif2ak3*/PERK fold change 0.96, P = 0.251). We emphasize that this cohort samples hippocampal rather than hypothalamic tissue and therefore provides only an indirect supraspinal snapshot of the trajectory of microglial and ER-stress transcriptional signatures after SCI; it is not equivalent to direct hypothalamic profiling at a later time point (see *Limitations*).

**Figure 4.**
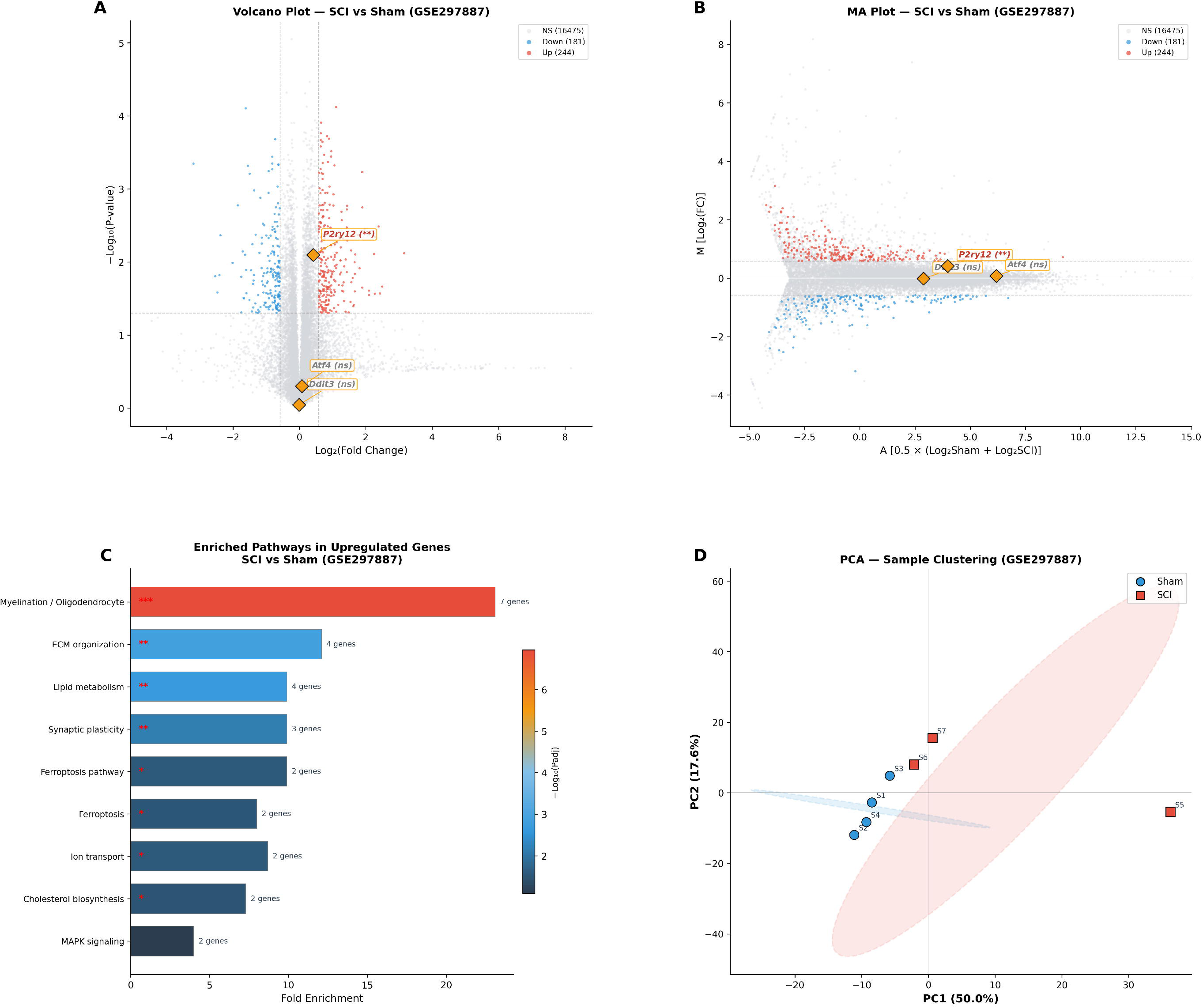
Independent validation of remote microglial activation with persistent ER stress silence in hippocampal regions following SCI (Validation Cohort: GSE297887, Chronic Phase). (A) Volcano plot of whole-transcriptome differential expression analysis (SCI vs. Sham) in remote hippocampal regions. Red: significantly upregulated genes (n = 244); blue: significantly downregulated genes (n = 181); gray: non-significant. Yellow diamonds highlight key markers: P2ry12 (significantly upregulated, **P = 0.008) and ER stress effectors Ddit3/CHOP and Atf4 (both non-significant). Dashed lines indicate significance thresholds (|FC| > 1.5, P < 0.05). (B) MA plot (Mean–Difference plot) providing an alternative visualization of differential expression as a function of average expression intensity. Key markers annotated as in (A). (C) Horizontal bar chart of enriched GO/KEGG pathways among upregulated genes, ranked by fold enrichment. The most prominently enriched pathway was Myelination/Oligodendrocyte (7 genes, Padj = 1.1 × 10⁻⁷), followed by ECM organization, lipid metabolism, synaptic plasticity, and—notably—ferroptosis (Alox12, Slc40a1; Padj = 0.028). The absence of canonical inflammatory pathway enrichment (NF-κB, TNF, TLR signaling) despite individual microglial gene activation suggests a non-classical ‘primed’ microglial state rather than overt neuroinflammation. *Padj < 0.05; **Padj < 0.01; ***Padj < 0.001. Fisher’s exact test with Benjamini–Hochberg correction. (D) Principal component analysis (PCA) of all 16,900 expressed genes. Blue circles: Sham samples (n = 4); red squares: SCI samples (n = 3). Shaded ellipses represent 95% confidence intervals. PC1 explains 49.6% and PC2 explains 17.6% of total variance, confirming substantial genome-wide transcriptomic separation between groups. Individual sample labels (S1–S7) are shown to illustrate inter-animal variability, with SCI sample S5 exhibiting the strongest transcriptomic shift along PC1.

## Discussion

By integrating a hypothesis-driven re-analysis of two independent transcriptomic datasets, we provide molecular evidence that the hypothalamic response to SCI unfolds as a temporally ordered cascade: an early, lesion-level-dependent neuroendocrine surge precedes — and is coupled with BBB disruption and early PERK-pathway sentinel-gene engagement — before the neuroinflammatory and ER-stress machinery becomes fully engaged. Below we discuss the principal observations, relate them to the broader literature, and explicitly address their limitations.

### A lesion-level-dependent neuroendocrine surge as a dominant early transcriptomic event

The most statistically robust transcriptional event in the post-SCI hypothalamus at day 7 was the coordinated upregulation of neuroendocrine genes (*Avp*, *Oxt*, *Ucn3*, *Asic4*), achieving genome-wide significance after stringent multiple-testing correction in the T3 SCI group but not the T10 SCI group. This molecular dichotomy parallels the clinical observation that dysnatremia after SCI is disproportionately associated with high-level lesions,^6,7^ and offers a transcriptomic counterpart to cellular stress-response programs described by Zhou et al. in the context of hypothalamic injury, where injury-induced activation of the ATF3/c-Jun/Lgals3 axis is associated with subsequent AVP-neuron loss and central DI.^30^ Our findings also complement and extend the multitissue framework of Zeng et al.,^22^ who originally generated the discovery dataset; their analysis identified T3-SCI-induced inflammatory hub genes and Gi-GPCR suppression at the network level, whereas our targeted gene-panel approach identifies the coordinated neuropeptide surge as the most individually prominent transcriptomic signal in the same dataset. The two analytical strategies are complementary rather than contradictory.

Mechanistically, the near-perfect AVP–OXT co-regulation (r = 0.941) suggests that both magnocellular systems are simultaneously engaged by the same non-osmotic drive. Because hypothalamic cells are particularly susceptible to ER-stress-induced dysfunction even under moderate stressor conditions,^31^ and because sustained or unresolved hypothalamic ER stress has been implicated in broad neuronal dysfunction,^32^ a prolonged surge of this magnitude can plausibly create pre-conditions for later ER proteostatic failure in a cell population whose vulnerability to ER stress is well documented.^17,18,33^

### Evidence of temporal ordering: absence of inflammation and ER stress at day 7

A key prediction of the cascade is that the neuroendocrine surge *precedes* the neuroinflammatory/ER-stress phase. Our data are consistent with this prediction: at day 7, classical neuroinflammation markers (*Il1b* P_adj_ = 1.00; *Gfap* P_adj_ = 0.905) and canonical UPR effectors (*Ddit3*/CHOP, *Atf4*, *Hspa5*/BiP, *Eif2ak3*/PERK) are all transcriptionally silent. This absence is consistent with the multi-step cascade by which SCI-induced hypothalamic inflammation is proposed to develop: initial spinal damage → systemic inflammatory mediator release → BBB compromise → peripheral immune infiltration. The observation that *Mmp9* is tightly correlated with *Avp* (r = 0.833; P = 0.0004) suggests that BBB disruption is already under way at day 7, running in parallel with the neuroendocrine surge, but that the downstream inflammatory consequences have not yet materialized in the parenchyma. Recent work on GSDMD-mediated inflammatory BBB breakdown^34^ and on MMP-9 as a central mediator of blood-spinal-cord-barrier breakdown after SCI^35^ is consistent with this interpretation. In systemic injury models such as myocardial infarction, ER stress and protein aggregation can be elicited in remote neural structures,^36^ establishing the principle that severe systemic injury can engage the UPR in remote neural tissue; our data suggest that at day 7 this ER-stress response has not yet become transcriptionally dominant in the hypothalamus. The biological bridge to subsequent ER stress is plausibly provided by microglia-derived IL-1β, which has been shown to engage the PERK/eIF2α/ATF4/CHOP axis in neurons,^37^ tying together the neuroinflammatory and ER-stress phases of the cascade.

### Sentinel genes: early PERK-pathway engagement without full UPR deployment

The coordinated sub-threshold shifts of Trib3 (fold change 5.9) and Ppp1r15a/GADD34 (fold change 2.5) provide exploratory clues that the earliest stages of PERK-pathway activation may have already commenced. TRIB3 is a direct ATF4–CHOP target,^25,38^ and has been identified as one of the most robust ATF4 target genes across multiple ChIP-seq and RNA-seq studies.^39^ This cohesive directional trend tentatively points to the functional engagement of ATF4 transcriptional activity, consistent with the known translational (rather than transcriptional) regulation of ATF4 protein by eIF2α phosphorylation.^40^ PERK–ATF4–CHOP axis activation has also been implicated in secondary injury and cell death across CNS injury modalities, including intracerebral hemorrhage.^41^ GADD34 provides an independent and complementary line of evidence: as the regulatory subunit of the PP1c phosphatase that dephosphorylates eIF2α, it serves as the canonical negative-feedback regulator of the PERK–eIF2α axis,^42^ and its upregulation indicates that PERK-mediated eIF2α phosphorylation has occurred. PERK-pathway engagement may not be limited to neurons: Zhu et al. demonstrated that PERK signaling in microglia governs the ER-stress–autophagy axis, and its dysregulation drives deleterious microglial activation after ischemic injury.^43^ The co-occurrence of oxidative stress indicators (*Mt1*, *Mt2a*) at the same time point further supports an ’early PERK engagement without full UPR deployment’ state, because oxidative stress is a well-established upstream trigger of the PERK pathway.^44^

### Cross-cohort temporal integration

Integrating findings across the two cohorts yields a coherent time-resolved snapshot: at day 7 (PRJNA953752), the hypothalamus shows maximal neuroendocrine output without significant inflammation or ER stress; in the chronic phase (GSE297887), remote supraspinal tissue (hippocampus) shows established microglial activation (*P2ry12* up) with continued ER-stress silence, reflecting the pan-encephalic penetration of the delayed inflammatory response. This pattern is consistent with a sequential cascade in which neuroinflammation develops with a delay of days to weeks after the initial neuroendocrine surge, while ER stress — predicted to peak transiently during the subacute window — is missed by both sampling windows. The shift in *P2ry12* from a non-significant downward trend at day 7 (fold change 0.52) to a significant upregulation in the chronic phase (fold change 1.33; P = 0.008) is compatible with a qualitative change in microglial state rather than a simple change in cell number. Microglial P2Y12 signaling is not confined to hemodynamic sensing: recent work has shown roles in calcium-dependent modulation of neurotransmission^45^ and in P2Y12-dependent modulation of central-nervous-system states.^46^ The causal relationship between hemodynamic disturbance, microglial activation, and remote brain injury is also supported by complementary vascular models: Kerkhofs et al. showed that pharmacological depletion of microglia and perivascular macrophages prevents vascular cognitive impairment in angiotensin-II-induced hypertension,^47^ establishing that microglial activation is mechanistically required for, not merely correlated with, hemodynamic-stress-induced brain damage. Recent studies further substantiate the concept of SCI-induced remote brain damage, including evidence that SCI-induced brain inflammation and cognitive decline can be reversed by spinal-cord cell transplants^48^ and that chronic SCI is associated with morphometric brain changes extending beyond sensorimotor networks.^49^

### Clinical implications and therapeutic window

From a translational perspective, the most important implication of our findings is the identification of a candidate therapeutic window between the early neuroendocrine surge and the later inflammatory/ER-stress cascade. At day 7, the hypothalamus is maximally engaged in compensatory neuropeptide production, but the destructive cascade has not yet fully developed. This window — tentatively spanning approximately post-injury days 3–10 on the basis of the integration of these two datasets — represents an opportunity for early intervention aimed at (i) reducing biosynthetic burden through judicious hemodynamic support (thereby reducing the non-osmotic drive for AVP hypersecretion); (ii) preserving BBB integrity through MMP-9 inhibition; (iii) pre-emptive ER-stress modulation using chemical chaperones such as tauroursodeoxycholic acid (TUDCA) or 4-phenylbutyric acid (4-PBA) and (iv) targeted microglial modulation and anti-ferroptotic strategies to mitigate established chronic neuropathology (**Supplementary Figure S1**). Consistent with this framing, we have reported in a propensity-matched retrospective cohort that dual-receptor therapy with pituitrin (AVP and OXT) reduced DI duration and mortality in patients with SCI.^50^ The present transcriptomic data offer one plausible mechanistic interpretation: exogenous AVP and OXT replacement may reduce the biosynthetic burden on endogenous magnocellular neurons, effectively unloading the ER and potentially delaying or preventing transition to ER-stress-mediated exhaustion. The near-perfect AVP–OXT co-regulation (r = 0.941) further supports dual-hormone replacement on the grounds that both systems are simultaneously stressed. We emphasize that this remains a mechanistic inference from transcriptomic data and retrospective clinical observations; prospective interventional testing is required before any such strategy can be recommended clinically.

### Limitations

Several important limitations must be acknowledged. First, both cohorts rely on publicly available datasets with relatively small sample sizes (n = 3–5 per group), limiting statistical power and likely contributing to the high P_adj_ values observed for many biologically plausible changes. The detection of sentinel genes such as *Trib3* and *Ppp1r15a* in the fold-change-filtered list despite non-significant P_adj_ values should be interpreted as exploratory findings requiring independent validation.

Second, the two cohorts differ in species (rat versus mouse), tissue specificity (hypothalamus versus hippocampus), injury model, and temporal sampling, precluding direct quantitative comparison. In particular, *the validation cohort samples hippocampal rather than hypothalamic tissue*, and the inferences drawn from it are necessarily indirect; hippocampal signatures cannot be assumed to recapitulate hypothalamic dynamics. These differences also serve, however, as a form of biological replication for features that do reproduce across species and tissues.

Third, bulk RNA-sequencing cannot resolve cell-type-specific contributions. The neuroendocrine surge signal likely originates from magnocellular neurons, whereas inflammatory and ER-stress signals may be diluted within the heterogeneous hypothalamic cell population. Single-nucleus RNA-sequencing or spatial transcriptomic studies will be essential to resolve these cell-type-specific dynamics.

Fourth, correlation analyses, while revealing strong associations (e.g., AVP–Mmp9 r = 0.833), cannot establish causality. The tight coupling between neuroendocrine and BBB genes may reflect shared upstream regulation rather than a direct mechanistic link. Conditional knockout models or targeted pharmacological interventions will be needed to establish causal relationships.

Fifth, both cohorts use male animals exclusively; generalization to female animals requires separate investigation. This is a substantive limitation given the known sexual dimorphism of neuroendocrine and neuroinflammatory responses.

Finally, and most critically, neither dataset captures the subacute window (approximately post-injury days 3–7 in the hypothalamic context) during which ER stress is predicted to peak. The present study therefore provides, rather than definitively demonstrates, a rationale for high-density temporal profiling (post-injury days 1, 3, 5, 7, 14) combined with hypothalamus-specific spatial transcriptomics to capture this critical but elusive molecular event.

## Conclusions

By integrating two independent transcriptomic datasets spanning the acute-to-chronic spectrum of SCI-induced supraspinal pathology, this study provides molecular evidence for a temporally ordered hypothalamic cascade after SCI. At post-injury day 7, the hypothalamic transcriptome is dominated by a massive, lesion-level-dependent neuroendocrine surge (*Avp* 7.2×, *Oxt* 14.3×, *Ucn3* 9.2×; all P_adj_ < 0.005), while classical neuroinflammation and ER-stress effectors remain transcriptionally silent. The concurrent detection of early PERK-pathway sentinel genes (*Trib3*, *Ppp1r15a*/GADD34), oxidative-stress indicators (*Mt1*, *Mt2a*), and a tightly coupled AVP–*Mmp9* BBB-disruption axis (r = 0.833; P = 0.0004) together define a molecular transition state poised at the threshold of the subsequent inflammatory/ER-stress phase. In an independent chronic-phase cohort, established microglial *P2ry12* activation with persistent ER-stress silence is consistent with delayed neuroinflammation and a transient ER-stress peak missed by both sampling windows. The neuroendocrine compensatory response therefore precedes, and may precipitate through biosynthetic overload and BBB compromise, the subsequent inflammatory/ER-stress assault hypothesized to drive AVP-neuronal exhaustion. The defined molecular window between early neuroendocrine activation and later inflammation/ER stress identifies a candidate therapeutic window that motivates future studies employing high-density temporal sampling and spatial transcriptomics to capture the elusive ER-stress peak and to define the precise timing of neuroprotective strategies in SCI.

## Transparency, Rigor, and Reproducibility Summary

### Study registration

This work was not pre-registered. It is a secondary, hypothesis-driven bioinformatic re-analysis of two publicly available transcriptomic datasets; the analytic plan — including the three pre-defined gene panels — was specified prior to interrogation of the full expression matrices and is described in full in the Materials and Methods.

### Sample size

Sample sizes were determined by the original depositors of each dataset. Discovery cohort (PRJNA953752): adult male Sprague-Dawley rats (sham, n = 3; T10 SCI, n = 5; T3 SCI, n = 5). Validation cohort (GSE297887): adult C57BL/6J mice (sham, n = 4; SCI, n = 3). No samples were added or removed. No a-priori power calculation was performed; given the fixed datasets, statistical power is limited and many biologically plausible effects fail to reach genome-wide significance (see Limitations).

### Inclusion / exclusion criteria

All samples available in each public deposit were included. For the validation cohort, genes with mean FPKM < 0.1 in both groups were excluded (yielding 16,900 expressed genes of 27,583 annotated) to avoid instability of log-fold-change estimates at very low expression.

### Blinding

Not applicable to data generation, as analyses were conducted on pre-generated, de-identified transcriptomic data. Sample-group labels were required for differential expression testing. The three hypothesis-driven gene panels (neuroendocrine, neuroinflammation, ER-stress/UPR) were pre-defined prior to interrogation of the data.

### Randomization

Not applicable; group assignments were set by the original experimental designs of the depositing investigators.

### Replication and validation

The primary DESeq2 analysis of the discovery cohort was independently verified with edgeR to confirm robustness of the core neuroendocrine findings. Cross-cohort concordance between the discovery (rat hypothalamus, day 7) and validation (mouse hippocampus, chronic phase) cohorts is discussed in the Results and Limitations; the two cohorts differ in species, tissue, and time point and are therefore not a direct technical replication.

### Statistical methods

Differential expression for the discovery cohort used DESeq2 (R 4.2.1) with Benjamini–Hochberg correction. Validation-cohort analyses used unpaired two-tailed Student’s *t* tests. Cross-sample correlations used Pearson’s r. Significance thresholds, pseudocounts, and filter criteria are detailed in Materials and Methods. No additional correction was applied for the *a priori* selection of the three gene panels, which is acknowledged as a limitation.

### Sex as a biological variable

Both cohorts use male animals exclusively (adult male Sprague-Dawley rats and adult male C57BL/6J mice per original depositions). Generalization to female animals requires separate investigation; this is acknowledged as a substantive limitation.

### Reagents and software

No wet-lab reagents were used. Software/packages: DESeq2 (Bioconductor), edgeR (Bioconductor), Python 3.12, NumPy, SciPy, pandas, scikit-learn, Matplotlib, seaborn. Specific version numbers and analysis scripts are available from the corresponding author on reasonable request.

### Data availability

All raw RNA-sequencing data are publicly available (NCBI Sequence Read Archive PRJNA953752; Gene Expression Omnibus GSE297887 / BioProject PRJNA1266676). All processed data and analysis scripts used to generate the figures and results are available from the corresponding author on reasonable request.

## Declarations

### Authors’ contributions

L.L. conceived the study, designed the bioinformatic analysis, performed the analyses, interpreted the results, and drafted the manuscript. H.Z. designed and performed the original animal experiments that generated the discovery cohort RNA-sequencing dataset (PRJNA953752) and contributed to data interpretation. J.G. contributed to data processing and clinical data interpretation. H.C. provided insights on clinical relevance. M.L. assisted with figure preparation and reference management. B.C. supervised the original transcriptomic data collection, provided critical review of the bioinformatic methodology, and contributed to manuscript revision. Z.L. supervised the project and provided critical intellectual input on neuroendocrine mechanisms. All authors read, critically revised, and approved the final manuscript. L.L. and H.Z. contributed equally to this work.

## Acknowledgments

We thank Professor Mou-wang Zhou (Peking University Third Hospital) and Professor Jin-wu Wang (Shanghai Ninth People’s Hospital) for facilitating access to the PRJNA953752 dataset and for their support of this collaborative analysis.

## Data availability statement

The RNA-sequencing data analyzed in this study are publicly available at the NCBI Sequence Read Archive (PRJNA953752; original publication: Zeng et al.^22^) and the Gene Expression Omnibus (GSE297887; BioProject PRJNA1266676). All analysis scripts and processed data files used to generate the figures and results are available from the corresponding author on reasonable request.

## Supplementary Material

**Supplementary Figure S1.**
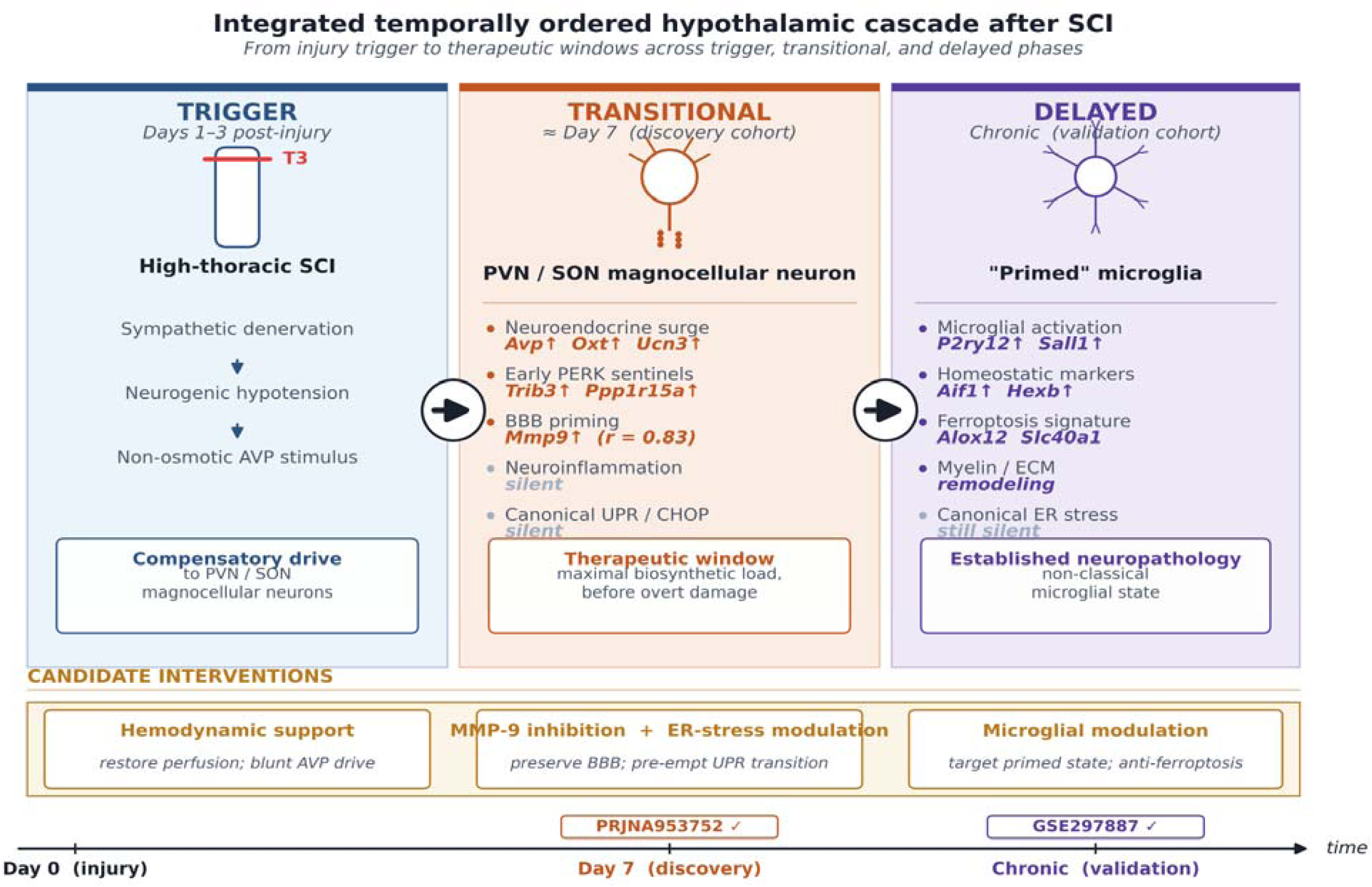
Integrated temporally ordered hypothalamic cascade after spinal cord injury. Integrated schematic organizing the proposed hypothalamic cascade into three sequential phases with their corresponding transcriptomic footprints and candidate therapeutic windows. **Trigger phase (days 1–3 post-injury):** high-thoracic (T3) SCI disrupts descending sympathetic outflow, producing neurogenic hypotension and a non-osmotic stimulus for magnocellular neurosecretion. **Transitional phase (≈ day 7, captured by the discovery cohort PRJNA953752):** PVN/SON magnocellular neurons exhibit a concurrent neuroendocrine surge (*Avp*, *Oxt*, *Ucn3*↑), early PERK-pathway sentinel engagement (*Trib3*, *Ppp1r15a*↑), and BBB priming through the tight AVP–*Mmp9* coupling (r = 0.83), while overt neuroinflammation and the canonical UPR/CHOP axis remain transcriptionally silent. **Delayed phase (chronic, captured by the validation cohort GSE297887):** a “primed” microglial signature emerges (*P2ry12*↑, *Sall1*↑) together with ferroptosis (*Alox12*, *Slc40a1*) and myelin/ECM remodeling signatures, whereas canonical ER-stress effectors remain silent. The transient ER-stress peak is hypothesized to occur within the subacute “temporal blind spot” between the two sampling windows. **Candidate interventions** (golden band) are mapped onto the phase in which they are predicted to be most effective: hemodynamic support to blunt the AVP drive during the trigger phase; combined MMP-9 inhibition and pre-emptive ER-stress modulation to preserve BBB integrity and abort the transition to the delayed phase; and microglial modulation (including anti-ferroptotic strategies) during the chronic phase. Coloured bullets indicate transcriptionally engaged events; grey bullets indicate events that are transcriptionally silent at the sampled time point. Arrows between columns denote the proposed causal sequence; the horizontal arrow at the base is the post-injury time axis with cohort badges marking the two sampling windows.

